# Secondary Growth and Exodermal Barriers Shape Local Root Hydraulics: Modeling Insights in Tomato

**DOI:** 10.64898/2026.01.27.701735

**Authors:** Marco D’Agostino, Rémy Schoppach, Adrien Heymans, Valentin Couvreur, Guillaume Lobet

**Affiliations:** Earth And Life Institute, UCLouvain, 1348 Louvain-La-Neuve, Belgium; Agrosphere IBG-3, Forschungszentrum Jülich, 52428 Jülich, Germany; Umeå Plant Science Centre, Department Of Forest Genetics And Plant Physiology, Swedish University Of Agricultural Sciences, Umeå, Sweden

**Keywords:** Root anatomy, hydraulic conductivity, secondary growth, modelling, tomato

## Abstract

Root water uptake efficiency depends on root system architecture and anatomical features of individual root segments. Beyond cell wall, membrane, and plasmodesmata hydraulic properties, root anatomy critically influences profiles of radial conductivity and axial conductance. While these structural factors have been well-characterized in monocotyledons, their role in dicotyledons—where developmental anatomy, secondary growth, and hydrophobic barrier dynamics differ—remains poorly understood.

Here, we integrate structural and functional models to assess how dicotyledon-specific anatomy, hydrophobic depositions (suberin/lignin in exo-/endodermis), and aquaporin contribution influence root hydraulics. Using tomato (*Solanum lycopersicum L*., cv. Moneymaker) as a dicotyledon model, our simulations show that:

- Exodermal suberin has negligible effects on radial conductivity when a lignin cap is present, and exodermal barriers are less effective than endodermal ones.
- Secondary growth and dicotyledon-specific anatomy are essential for sustaining high axial conductance, ensuring efficient water uptake across soil profiles and maintaining root system hydraulic conductance.

## Introduction

In response to the challenges imposed by changing climate conditions and the need for sustainable food production, there is a growing focus on the development of water-efficient and climate-robust crops (Tebaldi and Lobell, 2008). In that purpose, root systems are increasingly recognized as crucial breeding targets (De Dorlodot et al., 2007), as they mediate plant interactions with soil, water, and nutrients, key factors under changing climate conditions (Lynch, 1995; Uga, 2021). A comprehensive understanding of root system architecture and hydraulics, together referred to as root hydraulic architecture, is essential for identifying local strategies that optimize water uptake (Maurel and Nacry, 2020).

The efficiency of water uptake by plants is closely tied to their root system architectures and the anatomical features of individual root segments (Doussan, 1998; Heymans et al., 2020). The root architecture defines the position of roots in the soil, affecting their access to water and nutrients, as well as their connectivity to one another (Lobet et al., 2014). Root anatomy describes the geometry of root tissues from the cell scale. It strongly influences root hydraulic properties at the root segment scale, hereafter termed “local”, such as the radial conductivity (*k*_*r*_) and specific axial hydraulic conductance (*k*_*x*_) (Heymans et al., 2020; Lynch et al., 2021). It is worth reminding that the hydraulic “conductance” is an extensive property of a porous medium or membrane that describes the water flow rate generated by a unit gradient of water potential, while “conductivity” and “specific conductance” relate to the associated intensive properties, here after normalization by root segment outer surface and longitudinal dimension, respectively. Cell-scale features, such as the deposition of hydrophobic barriers (Wang et al., 2019) or aerenchyma formation (Fan et al., 2007) also have a strong influence on the local hydraulic properties. The integration of both root structural (architecture, anatomy) and functional (hydraulic conductivity) properties is needed to better estimate the water uptake capacities of specific plants (Passot et al., 2019).

To understand the latter, we first need to define water uptake in roots, particularly from the soil to the xylem. The radial movement of water through the cell anatomical network is commonly divided in two parallel pathways; the apoplastic pathway (through cell walls), and the cell-to-cell pathway (“transmembrane” across cell plasma membranes, and “symplastic” through plasmodesmata) (Steudle, 2000). Apoplastic flow can be blocked by the formation of hydrophobic barriers, with substances such as lignin (e.g. Casparian strips, lignin cap), while suberin hinders transmembrane flow (when forming waxy secondary walls covering the cell plasma membrane), in the endodermis and/or the exodermis (for reviews, see Enstone et al., 2002; Geldner, 2013). Furthermore, water flow in the cell-to-cell pathway is regulated by the aperture of plasmodesmata (see Bayer and Benitez-Alfonso, 2024 for review) and the permeability of cell membranes via aquaporins (Vandeleur et al., 2009; Couvreur et al., 2018). All of these cell-scale features and their interplay highly influence *k*_*r*_ (see Heymans, 2022 for sensitivity analysis).

Structural anatomical features also influence *k*_*r*_ and define *k*_*x*_. Notably, *k*_*r*_ is highly influenced by cortex width, or stele/cortex area ratio (Chimungu et al., 2014; Heymans et al., 2020), while *k*_*x*_ is dependent on the number and size of xylem vessels (Clément et al., 2022). It is worth mentioning that experimental measurements of *k*_*x*_ can deviates from simple computation of Hagen-Poiseuille equation because of finite xylem vessel lengths and increased resistance due to border pits (Choat et al., 2008; Boursiac et al., 2022).

These effects of structural anatomical features on *k*_*r*_ and *k*_*x*_ have been extensively studied in monocotyledons like maize, wheat or wheatgrass (Sanderson et al., 1988; Heymans et al., 2020; Heymans et al., 2021) and small dicotyledons like Arabidopsis or lupin (Meunier et al., 2018; Boursiac et al., 2022). However, there is still a lack of knowledge on the influence of specific anatomical traits on hydraulic properties in woody plants (like tomato or grapevine), where clear anatomical differences must impact root hydraulics (Strock and Lynch, 2020).

First, monocotyledons and dicotyledons root anatomies differ in terms of tissues and layers organization, most notably in the patterning of metaxylem and phloem (Lynch et al., 2021; Clément et al., 2022). In addition, dicot roots can undergo secondary growth (even in annual plants), which consist in a radial expansion due to the development of the stele, and an increase in parenchyma, phloem and xylem vessels, leading to an overall increasing diameter. This could have a significant effect on the radial hydraulic conductivity (Strock and Lynch, 2020).

Second, the timing and type of hydrophobic depositions can differ between dicotyledons and monocotyledons. In monocotyledons such as maize and wheat, the endodermis serves as the primary site for suberin and lignin deposition. However, in most angiosperms, including tomato but excluding Arabidopsis, apoplastic barriers also form within the exodermis. In tomato, recent advances in functional genetics unravelled the formation of suberin and lignin in exodermis. More specifically, Manzano et al., (2025) showed that in tomato, exodermis forms a polar lignin cap starting from 0.4 cm from the tip and with maximum lignification at 8 cm from the tip. Cantó-Pastor et al. (2024) showed that suberin deposition do not happen in endodermis, but well in exodermis, first as “patchy suberized” starting between 3 and 5 days, then as “continuously suberized” after 5 days. Both papers linked these apoplastic barrier depositions with water stress adaptation, by investigating the expression in different water conditions, or using tracers, respectively.

While this progress is substantial, the interaction between these hydrophobic depositions, secondary growth formation, and specific anatomical patterns must be studied to fully capture radial water dynamics in roots. However, such investigations are technically challenging experimentally (Passot et al., 2019; De Swaef et al., 2022).

Over the past decade, the scientific community has developed so-called S*tructural Functional Plant Models* (FSPM). *Structural* models represent the plant structure at different scales: root anatomy (e.g. Heymans et al., 2020; Sidhu et al., 2023), root system architecture (e.g. Pagès et al., 2014; Schnepf et al., 2018) or whole plant system (Zhou et al., 2020). These structural models are designed to be coupled to *functional* models, notably hydraulic models, to simulate water uptake at the anatomical scale (Couvreur et al., 2018), in a root system hydraulic architecture (Doussan, 1998; Meunier et al., 2020) or in the whole plant framework (Giraud et al., 2023). They have been used to investigate the interaction between root architectural and/or anatomical traits and water uptake (e.g. Zarebanadkouki et al., 2016; Meunier et al., 2018; Boursiac et al., 2022; Heymans, 2022; McLaughlin et al., 2024; Yu et al., 2024).

In this work, we used experimental data and computational models of roots - the cross-section structural generator GRANAR (Heymans et al., 2020) linked to the hydraulic solver MECHA (Couvreur et al., 2018) - to study how combined changes in root anatomy (including secondary growth), hydrophobic depositions levels (suberin and/or lignin in endo- and/or exodermis) and aquaporin contribution to membrane conductivity influence the root hydraulic properties in tomato (*Solanum lycoperiscum L*., var. Moneymaker), and to what extend it can shape hydraulic properties at root system scale.

## Results

### Quantification of anatomical features

We analysed anatomical features of 45 main root cross-sections from ten 30-day-old tomato plants (*Solanum lycopersicum L. var. Moneymaker*). From these observations, we classified tomato root anatomy into 3 phases: *maturation, intermediate* and *secondary growth*. Note that the meristematic and elongation zones were too short to be characterized by our methods (<5mm from the tip). Because roots did not have the same length, we choose to express the localisation of measurements in terms of days of growth instead of distance to tip.

The first phase is the *maturation* stage, which expanded from 0 to 5-7 days (picture A in upper panel of Figure 1). It was characterized by two blades of xylem across the stele. Xylem and phloem were organized in poles, with two blades of xylem sectioning the inner stele. The cortical cylinder was composed of one layer of endodermis, 3 to 4 cortex cell layers, one layer of exodermis and one layer of epidermis.

**Figure 1.**
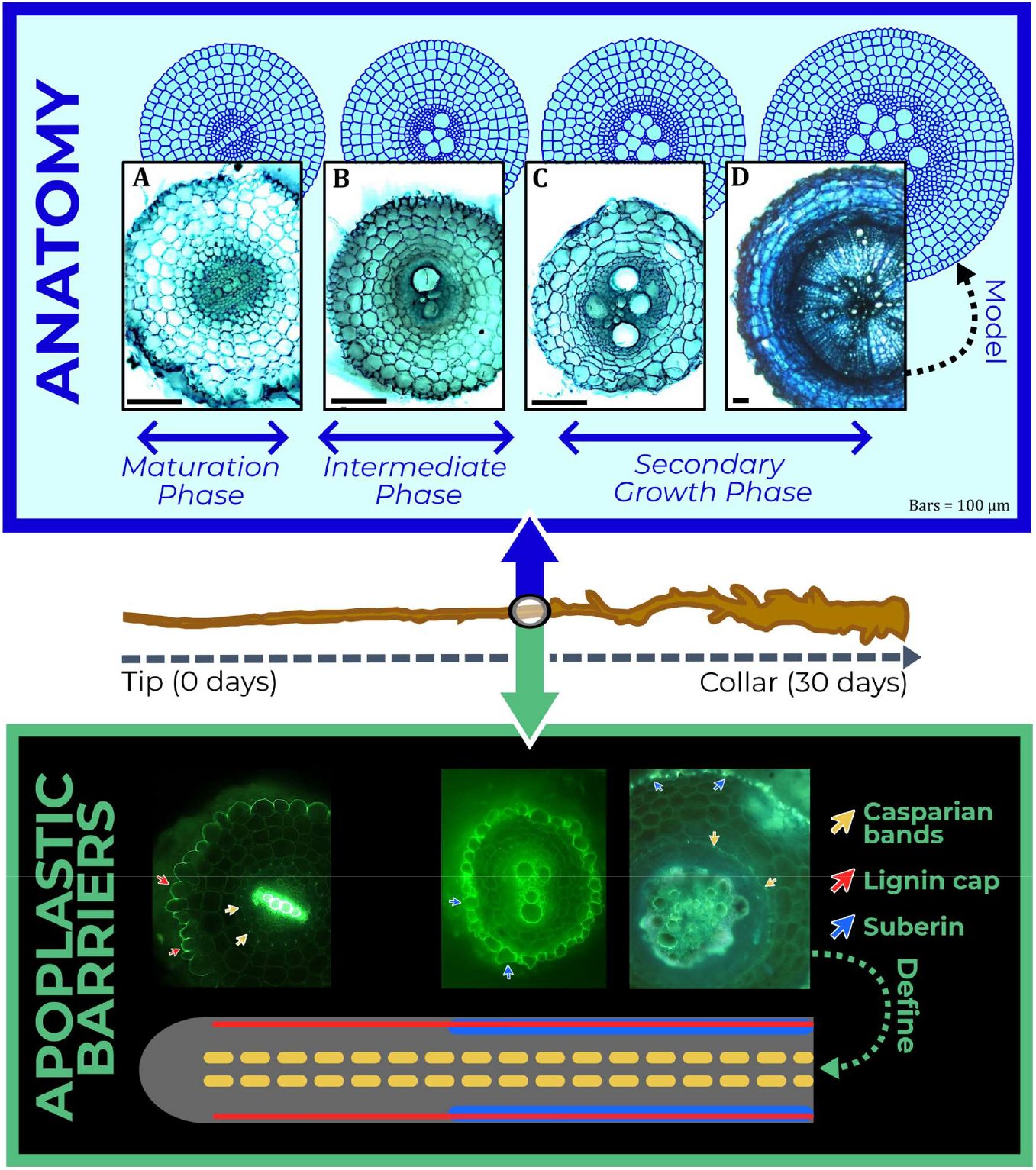
Quantification of tomato root anatomy features and hydrophobic barriers. Tomato roots (middle) were used to characterize and model anatomy (top) and hydrophobic barriers (bottom). The observed anatomy (top) was monitored and quantified to update and train a secondary growth model of the structural model GRANAR. The observed maturation (bottom) in terms of lignin and/or suberin formations in endodermis and exodermis served to build a map of the maturation timings.

The second phase, that we called *intermediate*, was characterized by the development of two new metaxylem files on each side of the xylem blades and expands from 5-7 to ∼25 days (picture B in upper panel of Figure 1). This stage was also characterized by an average of 3 cortex cell layers. At the end of this phase, the cambium/phloem began to form a ring around the inner stele, slightly introducing secondary growth.

Finally, the *secondary growth* stage (pictures C and D in Figure 1), starting approximately after 25 days, consisted of further exponential growth of the stele, with more parenchyma and xylem vessels, together with a large increase of xylem vessel size (Supplemental 1). Xylem patterning switched from blades to a heterogeneous disseminated placement in the inner stele (which is the main difference between pictures B and C in Figure 1). The phloem tissue multiplied, expanding into multiple layers, ranging from 2 to 5, while concurrently, the cortex was reduced to 2 layers (picture D in Figure 1).

The root radius along its longitudinal axis followed two trends: first it could be approximated as a constant (0.2 ± 0.03 *mm*) until the root segment age was 25 days (*P* < 10^-3^). After, it increased with age, so we built a linear regression of root radius as a function of root age with a Box-Cox transformation (*P* < 10^-4^; *R*^2^ = 0.729). This function was then used to predict the root radius of segments as a relation of their age.

Among the anatomical parameters, most of them showed a Pearson correlation coefficient (denoted *r*) equal or higher than 0.5 with the root segment age, except in in the cell diameter of phloem sieve tubes (*r* = 0.42) and cell diameter of pericycle (*r* = 0.35), which both displayed substantial variability regardless of root segment age. Notably, a negative correlation (*r* = −0.69) was spotted between the number of cortex layers and age, illustrating a reduction from 4 layers to 2 layers along the longitudinal axis of the root.

The strongest association between anatomical features and the age of the root segment was the diameter of xylem vessels (*r* = 0.96), the maximum size of xylem (*r* = 0.95) and the number of phloem layers (*r* = 0.92). In addition, a positive correlation between the root radius and root age was observed (*r* = 0.6) independently from the number of cortex layers (*r* = −0.29) and the proportion of phloem (*r* = 0.4). The root radius was best correlated to the whole stele diameter (*r* = 0.99), the diameter of stele parenchyma cells (*r* = 0.95) and the number of metaxylem vessels (*r* = 0.94).

### Discretization of hydrophobic deposition levels along the root axis

First, we observed the formation of Casparian strips in the endodermis starting at the maturation zone of the root, and the deposition of lignin in the outer-periclinal and anticlinal cell walls of the exodermis, forming a polarly localized lignin cap. We defined these two layers as deposition level *En*.*CS-Ex*.*LC* (for Endodermal Casparian Strips – Exodermal Lignin Cap). Around the age of 7-10 days, the exodermis showed a complete suberin deposition, while the endodermis still showed a simple Casparian strip; we defined this second deposition level as *En*.*CS-Ex*.*Sub* (for Endodermal Casparian Strips – Exodermal Suberization). The lower panel of Figure 1 summarizes the evolution of the different hydrophobic deposition levels.

### Development of a secondary growth module in GRANAR and estimation of local hydraulic properties

The updated version of GRANAR simulates root transverse anatomies that can present secondary growth. A complete description and the list of parameters are presented in Supplemental 2. The secondary growth module is a set of R functions that were directly integrated to the main code of GRANAR. With the updated version, users can choose to generate dicotyledon root cross-sections in primary growth stage or in secondary growth stage. Examples of simulated root cross-sections with secondary growth are shown in the top box of Figure 1. For the 40 quantified root anatomies, the measurements were translated into GRANAR parameters, and a virtual root anatomy was generated. Then, we used MECHA to compute *k*_*r*_ and *k*_*x*_ on each of the virtual anatomy, for 10 values of *kAQP* and 6 different hydrophobic deposition types (Table 1).

**Table 1.** Definition of hydrophobic depositions, for tomato and contrast comparison.

| *Type* | *Name* | *Description* |
| --- | --- | --- |
| Tomato (experimental) | ***En.CS*** | Endodermal Casparian strips |
|  | ***En.CS-Ex.LC*** | Endodermis Casparian strips and exodermal lignin cap |
|  | ***En.CS-Ex.Sub*** | Endodermis Casparian strips and exodermal suberization |
| Contrast Comparison (theoretical) | ***En.Sub*** | Endodermal suberization |
|  | ***En.Sub-Ex.CS*** | Endodermal suberization and exodermal Casparian strips |
|  | ***En.Sub-Ex.Sub*** | Endo and exodermal suberization |

### Investigation of the effect of anatomical features on radial conductivity

We first analysed the correlations between *k*_*r*_ and anatomical quantifications in the case of endodermal Casparian strips (*En*.*LC*) and mean *kAQP* value (4 × 10^-4^ *cm hPa*^-1^ *d*^-1^). *k*_*r*_ is negatively correlated (*r* ≤ −0.5) with most anatomical features except maximum xylem size, xylem cell diameter, phloem layers number, phloem proportion and cortex layers number. Notably, *k*_*r*_ is highly negativly correlated with cortex cell diameter (*r* = −0.76), radius (*r* = −0.75) and stele layer diameter (*r* = −0.65).

We made a linear mixed model to explain the variance of *k*_*r*_ by changes in age, radius, aquaporin contribution to membrane permeability, and hydrophobic deposition types. All variables have a significant effect on *k*_*r*_ (at *P* < 10^-3^). The generated linear model explains 92.4% of the variance of *k*_*r*_ and was used as predictive model for the next sections.

#### Specific effects of hydrophobic barriers deposition on radial conductivity

To assess the influence of hydrophobic barriers on *k*_*r*_, we performed a comparison of the mean of *k*_*r*_ while fixing aquaporin contribution at its average value (4 × 10^-4^ *cm d*^-1^*hPa*^-1^).

Our analysis showed that the presence of an exodermal lignin cap has no significant difference in terms of *k*_*r*_ compared to exodermal suberin (Figure 2). Even combined with either an endodermal Casparian strip or suberized endodermis, the resulting *k*_*r*_ values were not significantly different from one another.

**Figure 2.**
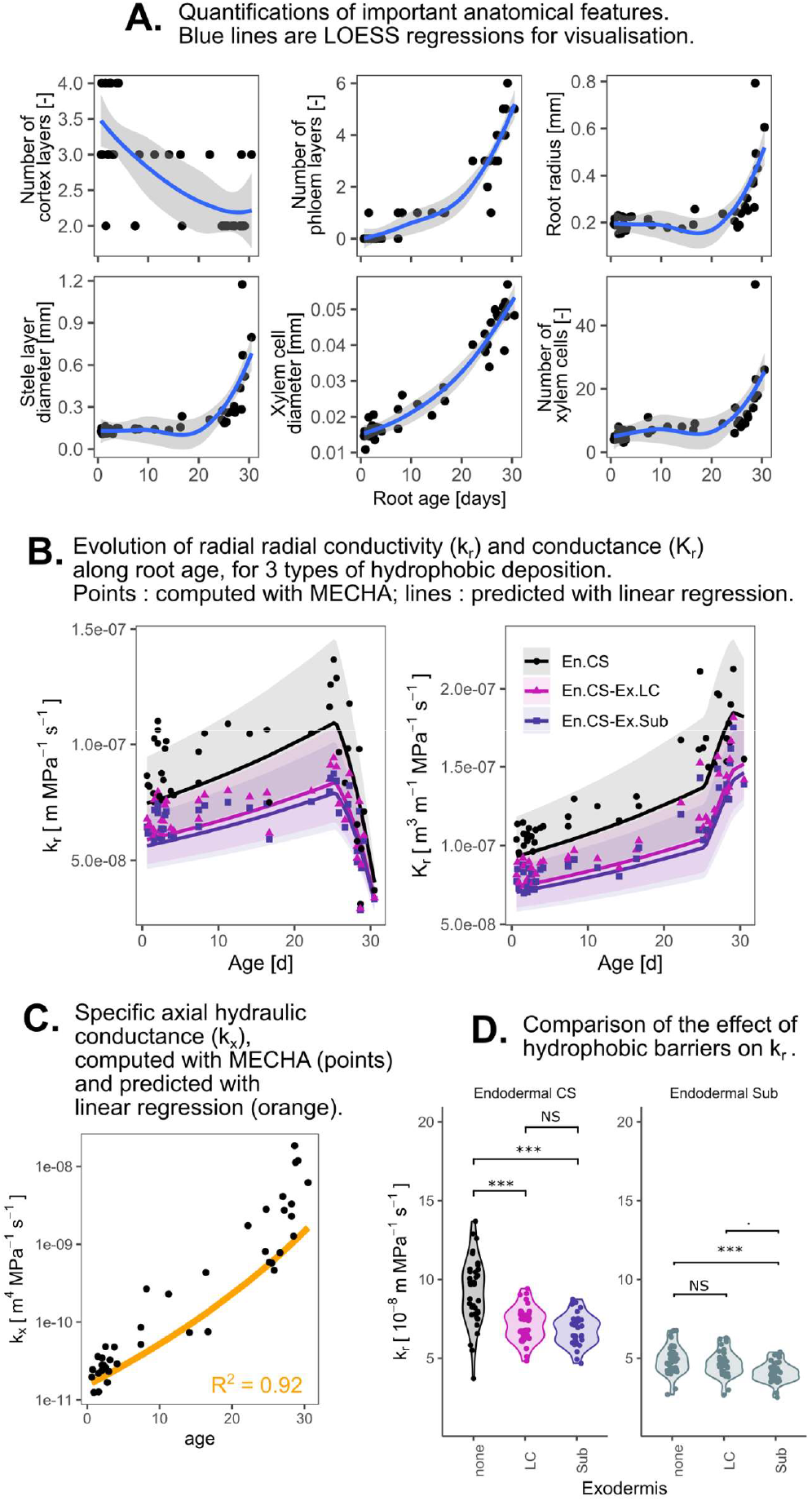
Quantitative analysis of root anatomical features, hydraulic properties, and the impact of hydrophobic depositions. uantifications of key anatomical features over root age. Data points represent measured values, while lines indicate LOESS ssions for visualization. (B) Evolution of radial conductivity (*k*_*r*_) and conductance (*K*_*r*_) along root age for three types of phobic deposition. (C) Specific axial hydraulic conductance (*k*_*x*_) computed with MECHA (points) and predicted with linear ssion (line). (D) Comparison of the effect of hydrophobic barriers on radial conductivity (*k*_*r*_) for endodermal casparian strips nd endodermal suberin (Sub).

In contrast, a clear reduction in conductivity emerged when an exodermal lignin cap was introduced relative to the case where there was no hydrophobic barrier in this cell layer. Specifically, adding a lignin cap to exodermal cells alongside an endodermal Casparian strip reduced *k*_*r*_ by 20.35%, whereas full suberization of the endodermis produced a stronger effect, lowering *k*_*r*_ by 45%, compared to a scenario with only an endodermal Casparian strip. When scaling these values to the level of the whole cross-section, radial conductance (*K*_*r*_) was calculated by multiplying the computed *k*_*r*_ of the cross-section by the perimeter. *K*_*r*_ then represents a flow of water per root length, per pressure unit and per time unit. Although the radial conductivity *k*_*r*_ drops (in average for the 3 deposition types) by 15.3% between 15-25 days and 25-30 days periods due to secondary growth, overall the radial conductance per unit length (*K*_*r*_) consistently increased by 18.9% between 15-25 days and 25-30 days because of the expanding root perimeter (Figure 2).

### Effect of radius and age on specific axial conductance

*k*_*x*_ is highly correlated with xylem vessel number and size due to the Hagen-Poiseuille relation (respectively *r* = 0.91 and *r* = 0.65) and is also highly correlated with root radius (*r* = 0.91). Since xylem cell diameter is highly correlated with root age (*r* = 0.96) and xylem cell number is highly correlated with root radius (*r* = 0.94), we constructed a linear model to predict *k*_*x*_ in function of root segment age and radius (*P* < 10^-3^; *R*^2^ = 0.92) (Table 3).

### Application: simulations of root system hydraulic architecture

To illustrate the effect of hydrophobic barriers and secondary growth on root system water uptake, we defined 4 *in silico* mutants; a wild type (*WT*), an aquaporin mutant (*aqp*), a suberin deficiency mutant (*sub*) and a combined secondary growth deficient – suberin deficient mutant (*sg-sub*) (Table 4). We assumed no interactions with other cell hydraulic or root anatomical traits.

We first generated 10 tomato-like simplistic root system architectures (RSA) of 60 days old with CRootBox (Schnepf et al., 2018). Then for each *in silico* mutant, *k*_*r*_ and *k*_*x*_ were computed for each root segment of the 10 RSA. It is worth noting that the axial conductance remains low for *sg-sub*, since there is no increase of diameter, compared to the WT and the other mutants.

For all *in silico* mutants, we computed the relative contribution of each segment to the total water uptake (*SUF*; Standard Uptake Fractions from Couvreur et al., 2012, under uniform soil water potential) and the root system hydraulic conductance *K*_*rs*_ [*m*^3^*MPa*^-1^*s*^-1^] with the root system hydraulic architecture solver MARSHAL (Meunier et al., 2020) every 10 days.

#### Root system hydraulic conductance

Statistical analyses showed no significant difference in *K*_*rs*_ between *aqp* and *sub* mutants compared to *WT* across all examined RSA ages for the selected cell-scale hydraulic properties (respectively *P* = 0.667 and *P* = 0.768 at 30 days-old and *P* = 0.021 and *P* = 0.253 at 60 days-old). In contrast, *sg-sub* mutant showed a reduction of *K*_*rs*_ compared to *WT* (40.3% at 30 days, 64.5% at 60 days, both at *P* < 10^-4^). A simple linear regression model incorporating RSA age, mutant type and their interaction effectively described *K*_*rs*_ variablility (*R*^2^ = 0.94). Under *P* < 0.05, the model showed no significant effect of *aqp* and *sub* compared to *WT*, and no significant effect of the interactions age-*sub*. By contrast, the effects of *sg-sub*, age, age-*aqp* interaction and age-*sg-sub* interaction were significative. These results show that the absence of secondary growth highly reduce *K*_*rs*_, and the the latter can be predicted linearly in function of the age.

#### Relative water uptake

No significant difference was detected in the whole water uptake profile between *WT* and *sg-sub* (under Kolmogorov-Smirnov test; *P* = 0.952 at 30 days old an *P* = 0.994 at 60 days old). Despite the absence of global differences, local shifts of relative water uptake at different soil layers were observed between *WT* and *sg-sub* at specific soil layers, at significance level of *P* < 5 × 10^-3^. For 30 days-old RSA, the relative water uptake for *sg-sub* is higher than for *WT* between 0 and 4 cm (+51.9% at 0-2cm; + 34.8% at 2-4cm depth), and is lower than *WT* between 12 and 14 cm (-11.3%). For 60 days-old RSA, the relative water uptake for *sg-sub* is higher than *WT* between 0 and 6 cm (+203.2% at 0-2cm; +123.1% at 2-4cm; +63.9% at 4-6cm), and is lower than *WT* between 12 and 22 cm (-33.3% at 12-14cm; -36.7% at 14-16cm; -30.2% at 16-18cm; -17.2% at 18-20cm; -8.1% at 20-22cm) then higher again between 24 and 28 cm (+7.4% at 24-26cm, + 9.5% at 26-28cm).

Altogether, these results show that, despite no apparent difference in whole root water uptake distribution, RSA without secondary growth (*sg-sub*) takes more water near the collar than the RSA with secondary growth, but takes less water in deeper soil layers.

## Discussion

### Anatomy and secondary growth effects on hydraulic properties

Radial conductivity is fundamentally determined by anatomical traits, such as the number of cortex cells and the structure of the stele, explicitly modelled in frameworks like MECHA (Couvreur et al., 2018). However, these traits are time consuming to measure experimentally. Our results indicate that root age and radius account for a considerable proportion of the observed variability in radial hydraulic conductivity (*k*_*r*_). Both also show a good correlation with anatomical traits, justifying their use as proxies for predicting *k*_*r*_. This aligns with observations reported by Heymans (2022).

With age, the number of stele parenchyma cells increases, as well as the number and the size of metaxylem vessels. The cambium generates secondary phloem and parenchyma cells, which increase the total size of the stele. The number of cortex cell layers decreases with time (from 3-4 in primary growth to 2 in secondary growth) and is not correlated with the increase of diameter. This reduction of cortex, at the opposite of what is observed in maize for example (see Chimungu et al., 2014; Heymans et al., 2020), contributes to an increase in *k*_*r*_ by reducing the length of the pathway for water to reach xylem vessels.

On the other hand, the number of cell layers of stele and phloem/cambium increases with age, especially in secondary growth zone. This increase makes them major contributors to the total increase in the root radius. With the increase of the stele comes an increase of xylem vessel number and cell size, which makes the specific axial hydraulic conductance *k*_*x*_ increase exponentially during formation of secondary growth. The increase of phloem and cambium layers eventually counters the effect of the thinner cortex layer and makes *k*_*r*_ drop.

### Influence of apoplastic barriers deposition levels on root radial conductivity

#### Additional suberin does not reduce *k*_*r*_ compared to lignin cap, in exodermis

Manzano et al. (2025) showed that in tomato, young parts of the roots present a lignin cap in the exodermis instead of Casparian strips. Cantó-Pastor et al. (2024) showed that tomato roots present suberization in exodermis but not in endodermis in 7 days old seedlings, which we also observed. We confirmed such observations through fluorescence microscopy, and we observed endodermal Casparian strip formation at the same timing.

Under the assumption that lignin is completely impermeable (in MECHA we set the specific hydraulic conductivities values for cell walls with suberin and/or lignin to zero), our simulations suggest that an exodermal lignin cap strongly reduces *k*_*r*_ by 20% compared to only endodermal Casparian strips. However, when a lignin cap is present, the impact of additional exodermal suberin on *k*_*r*_ is so small that it is overshadowed by the effect of the natural variability in root anatomies. Our interpretation is that the lignin cap is sufficient to force water flow through the symplastic pathway, so adding an additional hydrophobic deposition on the opposite face of the exodermis has no substantial effect. However, our approach does not include changes of other hydraulic parameters linked with hydrophobic deposition, like aquaporin expression (Hachez et al., 2006; Calvo-Polanco et al., 2021), which may play a significant role in regulating radial conductivity.

It is also worth noting that our approach only consider the role of suberin as a barrier for the transmembrane and apoplastic transport of water, and does consider its other roles, like the preservation of homeostasis and nutrient transport, a response to various biotic/abiotic stresses, pathogen resistance or bacteria community composition (Andersen et al., 2015; Barberon et al., 2016; Salas-González et al., 2021; Liu and Kreszies, 2023).

#### *k*_*r*_ is more sensitive to suberization in endodermis compared to exodermis

Our results show that in tomato, suberization would have a stronger impact on *k*_*r*_ if it were to happen in the endodermis rather than the exodermis. Adding a suberized endodermis makes *k*_*r*_ decrease by 46% compared to a simple endodermal Casparian strip, while adding and exodermal barrier (either lignin cap or suberin) to endodermal Casparian strip makes *k*_*r*_ decrease by 20% compared to only endodermal Casparian strips (Figure 2).

An explanation is that forcing a fully symplastic pathway across the endodermis is more of a bottleneck due to its smaller surface relative to the exodermis. In addition, in default MECHA inputs, exodermis cells present a higher plasmodesmata frequency compared to endodermis cells. These values were originally derived from maize (*Zea mais L*.), for which MECHA was initially parameterized (Clarkson et al., 1987). They were kept for tomato due to the absence of species-specific data. Given that the exodermis not only possesses a higher plasmodesmata frequency but also a greater surface area relative to the endodermis, the symplastic pathway is more conductive in the exodermis than in the endodermis. Consequently, suberization forcing a fully sympastic pathway provoked a more reduced *k*_*r*_ when happening in the endodermis than in the exodermis.

### Influence of aquaporins on root radial hydraulic conductivity

The contribution of aquaporin to membrane conductivity (*kAQP*) influences the root radial conductivity, depending on the deposition level (Figure 2 and Table 2). The main hypothesis for that is that with no suberized layer, aquaporins enables the switch from apoplastic pathway to symplastic pathway through the transmembrane pathway. Once the exodermis (or endodermis) is suberized, water can only pass across through the symplastic pathway that involves a combination of transport in series across membranes and plasmodesmata. The latter become the primary limit to radial water transport, and therefore reduce the sensitivity of radial conductivity to aquaporins (Gambetta et al., 2013; Heymans, 2022). In that sense, for example, Gambetta et al. (2013) showed that aquaporin expression in grapevine roots decreased in suberized secondary growth part compared to un-suberized, younger zones.

**Table 2.** Summary of the linear regression of radial conductivity in function of hydrophobic barriers, aquaporin contribution, age and radius.

| *Variable* | *Estimate* | *Std. Error* | *t value* | *P value* |
| --- | --- | --- | --- | --- |
| (Intercept) | -2326 | 14.14 | -164.6 | $< 10^{-4}$ |
| age | 15.33 | 0.2718 | 56.4 | $< 10^{-4}$ |
| radius | -2307 | 24.54 | -94.01 | $< 10^{-4}$ |
| *En.CS-Ex.LC* | -264.5 | 18.7 | -14.14 | $< 10^{-4}$ |
| *En.CS-Ex.Sub* | -302 | 18.7 | -16.15 | $< 10^{-4}$ |
| *En.Sub* | -454.7 | 18.7 | -24.31 | $< 10^{-4}$ |
| *En.Sub-Ex.CS* | -581.5 | 18.7 | -31.09 | $< 10^{-4}$ |
| *En.Sub-Ex.Sub* | -701.3 | 18.7 | -37.5 | $< 10^{-4}$ |
| kAQP | 718616 | 21314 | 33.72 | $< 10^{-4}$ |
| *En.CS-Ex.LC* : kAQP | 10200 | 30142 | 0.3384 | 0.7351 |
| *En.CS-Ex.Sub* : kAQP | -33898 | 30142 | -1.125 | 0.2609 |
| *En.Sub* : kAQP | -409658 | 30142 | -13.59 | $< 10^{-4}$ |
| *En.Sub-Ex.CS* : kAQP | -298560 | 30142 | -9.905 | $< 10^{-4}$ |
| *En.Sub-Ex.Sub* : kAQP | -378800 | 30142 | -12.57 | $< 10^{-4}$ |

Adjusted $R^2 = 0.924$

**Table 3.** Summary of the linear regression of specific axial conductance in function of age and radius.

| *Variable* | *Estimate* | *Std. Error* | *t value* | *P value* |
| --- | --- | --- | --- | --- |
| (Intercept) | -231.9857 | 5.0813 | -45.655 | < 10-4 |
| radius | 53.7887 | 22.3322 | 2.409 | 2.10E-02 |
| age | 3.7873 | 0.2473 | 15.313 | < 10-4 |

Adjusted R<sup>2</sup> = 0.919

**Table 4.** Definition of the *in silico* mutants.

| Name | Endodermal deposition | Exodermal deposition | Secondary growth | Aquaporin contribution |
| --- | --- | --- | --- | --- |
| *WT* | Casparian strips | Lignin cap and suberization | Yes | Constant |
| *aqp* | Casparian strips | Lignin cap and suberization | Yes | **Decreasing** |
| *sub* | Casparian strips | **Lignin cap** | Yes | Constant |
| *sg-sub* | Casparian strips | **Lignin cap** | **No** | Constant |

### Application to root system hydraulic architectures: root system conductance and standard uptake fractions

Our results show that root system hydraulic conductance *K*_*rs*_ is highly influenced by secondary growth (Figure 3). The explanation is that secondary growth creates a much higher axial conductance in upper parts of the root (due to higher xylem vessels size and number), which enables the root system to transport more water and therefore take up more water from deeper soil layers. Indeed, the profile of the standard uptake fractions (*SUF*, which are the relative contribution of each segment to the total water uptake in homogenous soil) of the RSA without secondary growth shows higher relative water uptake near the collar, indicating that the root segments near the collar still take up water. But since the axial transport is limiting without secondary growth, one could expect that secondary growth root systems are able to take more water from deepest soil layers.

**Figure 3.**
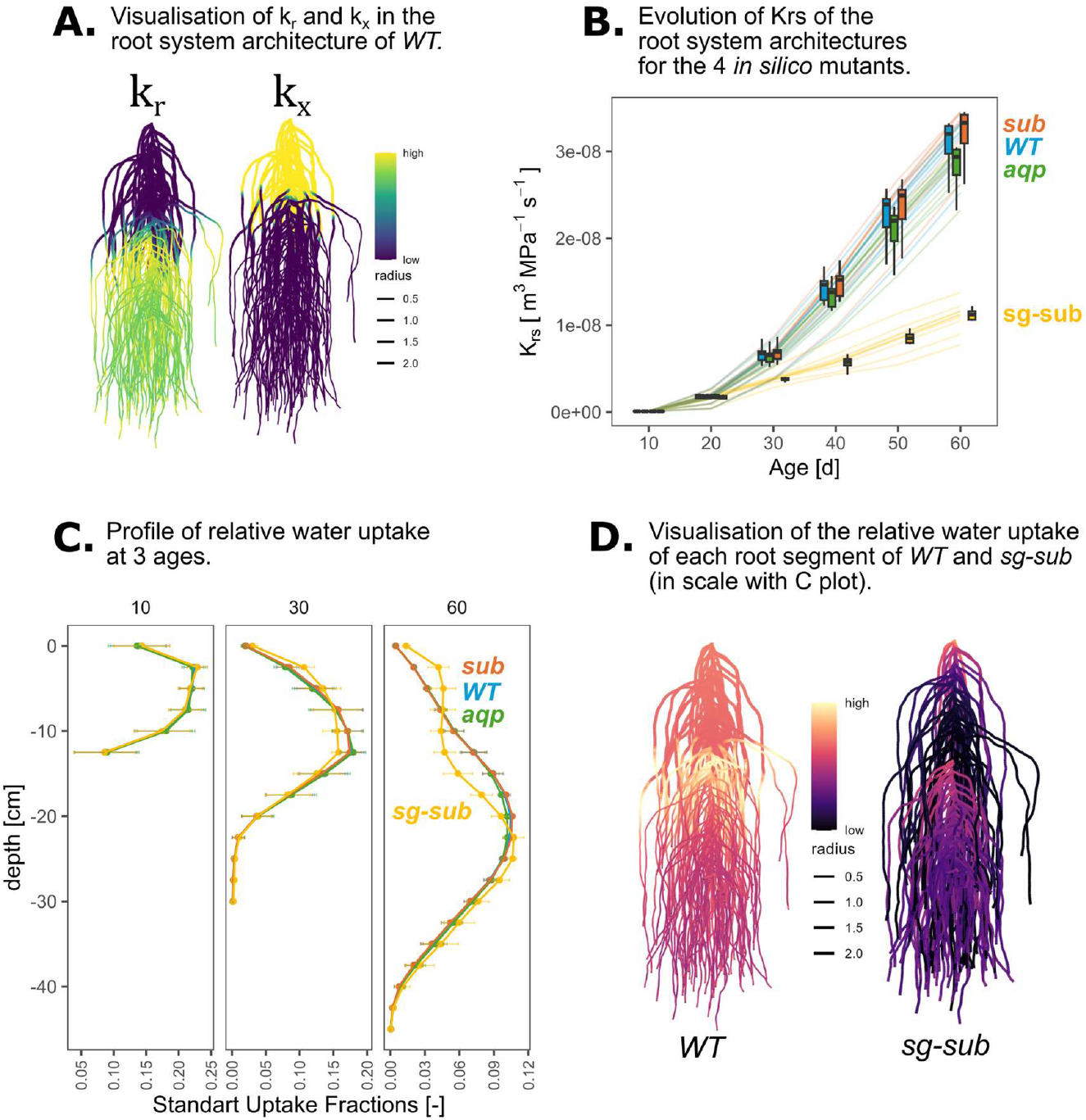
Visualization and analysis of root system architectures and water uptake profiles in wild-type (*WT*) and in silico mutant plants. (A) Visualization of radial (kr) and axial (kx) hydraulic conductivities in the root system architecture of WT plants. The color gradient represents root radius (high to low). (B) Evolution of root system hydraulic conductivity (*K*_*rs*_) over time for the four in silico mutants. (C) Profile of relative water uptake at three developmental ages (10, 30, and 60 days). Standard uptake fractions are plotted against depth for *sub, WT, aqp*, and *sg-sub* mutants, highlighting differences in water acquisition strategies over time. (D) Visualization of the relative water uptake of each root segment in *WT* and *sg-sub* mutants. The color gradient indicates water uptake intensity.

Another point is that, with secondary growth comes higher external surface of the root, which makes the radial conductance *K*_*r*_ increase, even if the radial conductivity *k*_*r*_ decreases. That means that secondary growth portions are still able to take up water radially, even with a much lower *k*_*r*_. This is why the relative water uptake (represented by *SUF*) does not decrease directly with the appearance of secondary growth. However, the relative water uptake ultimately collapses near the collar for two main reasons: there are less roots at this point in the profile, and the large axial conductance facilitates the delocalization of water uptake to other parts of the root system. This highlights the functional importance of the ratio between radial and axial conductivity (Draye et al., 2010).

### Opening to future works

In this work we developed a framework to couple plant hydraulic models from cells scale to root system scale, to integrate local knowledge into upscaled scale, similarly to what was developed in (Heymans, 2022) but for dicotyledons. Our sensitivity analysis included aquaporins, exodermis apoplastic barriers and secondary growth.

All our hydraulic scenarios include a lignin cap in the exodermis, which appears in the first 2 cm of the root tip (Manzano et al., 2025). Our results show that that lignin cap has a strong effect on *k*_*r*_, and therefore limits it from the very tip of the root. Our simulations also confirm that aquaporins highly influence transmembrane transport. However, we did not include in our sensitivity analysis the effect of plasmodesmata, which control symplastic water transport and could lead to significant changes in hydraulic properties. Unfortunately, data about plasmodesmata conductance in tomato is lacking. Further work can be done to investigate the effect of plasmodesmata and include it into further sensitivity analysis.

Our framework enables quantitative assessment of root system hydraulic performance under varying environmental and physiological conditions. Its modular design supports the integration of novel hypotheses, such as comparative analyses of water-use efficiency in diverse root architectures (Chandrasekhar and Julkowska, 2022). Furthermore, the pipeline allows systematic evaluation of how alterations in hydraulic traits—whether structural (e.g., xylem anatomy, tissue composition) or functional (e.g., membrane permeability, apoplastic barriers)— influence water acquisition, facilitating future studies on mutants with modified hydraulic properties (Cantó-Pastor et al., 2024; Manzano et al., 2025).

In this study, we have established a framework that integrates plant hydraulic models from the cellular level to the root system scale, enabling the incorporation of local insights into broader-scale analyses, akin to the approach done in (Heymans, 2022) but tailored for dicotyledons. Our sensitivity analysis included factors such as aquaporins, exodermis, apoplastic barriers, and secondary growth. Future investigations using this framework could explore additional parameters, including plasmodesmata, and assess various combinations, such as different timings of lignin cap deposition.

## Conclusion

In this work, we investigated the effect of the interplay between root developmental anatomy, secondary growth and hydrophobic deposition level on root hydraulic properties. We used and updated existing process-based computational models, and we integrated experimental root data and information on root system scale modelling.

Our results show that the exodermal lignin cap significantly reduces radial conductivity, but an additional layer of suberin in the exodermis does not when accounting for natural variability in root anatomies, thus questioning the role of suberin as hydrophobic barrier in exodermis.

Our simulations also show that secondary growth compensates for the decrease of conductivity by increasing the absorption surface, eventually making radial hydraulic conductance increase with the root age. Like so, secondary growth would compensate the effects of hydrophobic barriers in radial uptake, in addition to exponentially increase the axial conductance, allowing for water from deepest root zones to move to the leaves. This is likely to be a critical feature in dicotyledons, where the totally of the water taken up by the root system must be funnelled through the root collar.

With our methodology, we confirmed experimental findings on the importance of exodermal hydrophobic depositions, and we created a modelling pipeline to simulate different scenarios of hydraulic properties, for example to investigate the hydrodynamics of apoplastic barriers mutants.

The models used in this study are purely process-based, user friendly and open source. The coupling and updates of such different multiscale functional structural plant models open avenues to improve and expand applications of digital avatars and studies on root system hydraulic dynamics.

## Materials & methods

### Plant growth & sample preparation

40 tomato (*Solanum lycopersicum* var. Moneymaker) plants were grown in UCLouvain greenhouses (Louvain-la-Neuve, Belgium) from October 12 to November 11, 2022, in an aeroponic system at 24°C. Seeds were directly dropped in small PVC tubes composed of a plastic web as a floor and filled with a mix of perlite and vermiculite (1:1) to allow the plant to stabilize. The tubes were fixed in expanded polyester plates (10x50cm) and mounted on top of the aeroponic system (Dorlodot et al., 2005). Watering was done by spraying Hoagland nutrient solution for 5 seconds every 3 minutes. Plants were collected after 30 days of growth. We selected the 9 healthier plants (that presented secondary growth) and harvested their root systems. Tap roots were cleaned from lateral roots and stored in WEG (water-ethanol-glycerol 3:3:2) as recommended by Kitin et al. (2020). Roots were divided into 1-3 cm long segments in which we considered anatomy as consistent, to replicate microscopy observations. The length of each segment of each plant as well as the length of the whole root were carefully measured. Based on previous manipulations and data, we simplified the root growth rate by simply taking the mean through the entire growth period. Since all roots were not the same length but the same age, we calculated the length/age ratio for each root. The position of each segment was then expressed in days instead of centimetres.

### Microscopy

Root segments were deposited in small moulds and embedded in a 40°C, 0.5 agar solution, following the Atkinson and Wells (2017) protocol. Once the agar solution became solid, cross-sections were made using a razor blade, and stored in Petri dishes in a drop of water. Then cross-sections were stained with a drop of Toluidine O and observed under microscope (Eclipse E400, NIKON). For each cross-section of each root segment, pictures were taken (Moticam 2000, MOTIC). For 3 main roots, additional staining was applied to reveal apoplastic barriers in each segment. Berberine Aniline was used to reveal Casparian strips (CS), while Fluorol Yellow was used to monitor suberized tissues, using protocols of Lux et al. (2005) and Kitin et al. (2020). For both colorations, microscopical pictures were taken under UV-excitation with an Eclipse E400 (NIKON).

### Anatomical features quantification

Cross-sections pictures were quantified using ImageJ (Schneider et al., 2012) to get data in the form of GRANAR parameters (Heymans et al., 2020). 45 cross-sections were quantified from the 9 different tap roots. The full quantification protocol consists of drawing a few cells for each tissue and measuring their size (in µ*m*^2^) or their width (in µ*m*). For metaxylem, each vessel cell was drawn, because the number of metaxylem vessels is a parameter of GRANAR, and it is obtained by counting the number of measurements. Protoxylem was considered as parenchyma. For parenchyma, phloem and cortex, cells were drawn along one or more radius, to avoid unintentional selection bias. For tissues that are composed of one single cell layer (pericycle, endodermis, exodermis, and epidermis), a representative subset of cells was drawn along the arc of circle of the cell layer. For phloem, pericycle, endodermis, cortex, exodermis and epidermis, the cell width was measured, oriented in the cross-section radius direction.

In secondary growth cross-sections, cambium was classified as phloem sieve tubes. Indeed, it was hard to distinguish the two tissues in our pictures, and we do not aim to do a developmental analysis. We therefore made the hypothesis that phloem and cambium have a similar behaviour in terms of radial hydraulic conductivity. Even if we expect greater axial conductance in phloem than in cambium, we hypothesize that it does not have a significant influence due to the relatively small proportion of cambium compared to phloem and xylem.

GRANAR generates round cells, whereas cells in these tissues appear to be elongated. We therefore chose to generate round cells in GRANAR but with a diameter equal to the width of the measured cells. By doing so, we end up with generated tissues that have more cells (that are round) in total, but with a diameter equal to the width of the measured cells in the direction of the cross-section radius. This choice was motivated by the fact that layer width has tremendous influence on radial water flow while the number of cells per cell layer (and by extension the number of cell walls for apoplastic pathway) have little effect (Heymans et al., 2020). All measurements of all pictures were stored in a database and treated in R (R Core Team, 2022) to get a table of the quantifications as GRANAR input parameters per plant and per segment.

### Update of GRANAR to generate secondary growth

The codes of GRANAR and the description of the workflow are available on GitHub (https://github.com/granar/granar).

It is worth noting that GRANAR present some limitations due to the nature of its implementation. Notably, the switch from primary growth to secondary growth is not smooth and corresponds to two different code workflows. While in primary growth mode GRANAR explicitly generates xylem blades, in secondary growth mode GRANAR generates randomly placed metaxylem vessels around the centre of the stele, then directly losing the typical blade formation.

### Computing water flow in hydraulic anatomies

We computed *k*_*r*_ and *k*_*x*_ of every quantified anatomy with MECHA (Couvreur et al., 2018) for 60 scenarios : 6 different deposition levels, and 10 values of contribution of aquaporin to membrane permeability *kAQP* [*cm d*^-1^*hPa*^-1^]. The other hydraulic parameters of MECHA were set as default values, coming from literature (Supplemental 4) (see Couvreur et al., 2018).

The different hydrophobic deposition levels are summarized in Table 1. This choice was supported by our own observations, as well as results from Cantó-Pastor et al. (2024) and Manzano et al. (2025). Monocot deposition levels are the standard ones for maize from Doussan (1998) and Heymans et al. (2020). The test values of *kAQP* range from 1E-04 and 1E-03 cm d^-1^ hPa^-1,^ to basically explore its influence around the mean value from maize; 4.3E-04 cm d^-1^ hPa^-1^ (Ehlert et al., 2009; Couvreur et al., 2018). It is useful to note that such units [cm d^-1^ hPa^-1^] were chosen for ease of reading instead of standard [m s^-1^ MPa^-1^].

### Statistical analysis of the influence of anatomical features on hydraulic properties

#### Effect of age, radius, aquaporins and hydrophobic barriers on radial conductivity

A Box-Cox transformation (Box and Cox, 1964) of parameter *λ* = −0.4242 was applied to *k*_*r*_ values, and improved the normality of the data, going from *P* < 2.2 × 10^-1^ to *P* = 1.97 × 10^-10^ under Shapiro-Wilk test. While the Shapiro-Wilk test statistic remained significant, the high explanatory power of the model, combined with visual validation of predicted versus observed values, supports the robustness of our approach.

#### Effect of age and radius on specific axial conductance

A Box-Cox transformation of parameter *λ* = −0.141 was applied to *k*_*x*_ data, to force normality; going from *P* = 0.001574 using neperian logarithm transformation to *P* = 0.00315 using Box-Cox (under Shapiro-Wilk test).

The linear regression model has the following equation:

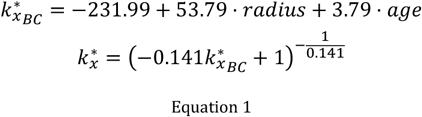

where 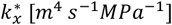 is the estimated specific axial conductance, after transformation with Box-Cox technique.

### Root system hydraulic architectures

10 virtual root systems were simulated using the CRootBox shiny app (plantmodelling.shinyapps.io/shinyRootBox/) (Schnepf et al., 2018). We used the *Arabidopsis thaliana* preset and modified some parameters based on our own data (see Supplemental 3), and by setting the deviation of these parameters to 30%. *Tropism parameter 1* was set to 1.5 for lateral roots (type 2).

We defined 4 *in silico* mutants by changing radius, *k*_*r*_ and *k*_*x*_ and s of each RSA segments (Table 4).

- *WT* : an *in silico* wild type of tomato.
  ∘ Hydrophobic barriers set successively as *En*.*CS-Ex*.*LC* than *Ex*.*CS-Ex*.*Sub*.
  ∘ Root radius of each segment applied as a function of the root segment age.
  ∘ *kAQP* set constant at 5 × 10^-4^ *cm* ⋅ *HPa*^-1^ ⋅ *d*^-1^.
  ∘ *k*_*r*_ and *k*_*x*_ of each segment calculated using the regression functions.
- *aqp* : an *in silico* mutant of aquaporins.
  ∘ Hydrophobic barriers set successively as *En*.*CS-Ex*.*LC* than *Ex*.*CS-Ex*.*Sub*.
  ∘ Root radius of each segment applied as a function of the root segment age.
  ∘ *kAQP* set as linearly decreasing from 5 × 10^-4^ to 1 × 10^-16^ *cm* ⋅ *HPa*^-1^ ⋅*d*^-1^ along root age.
  ∘ *k*_*r*_ and *k*_*x*_ of each segment calculated using the regression functions.
- *sub* : an *in silico* mutant of suberin.
  ∘ Hydrophobic barriers fixed to *En*.*CS-Ex*.*LC* for all root segments.
  ∘ Root radius of each segment applied as a function of the root segment age.
  ∘ *kAQP* set constant at 5 × 10^-4^ *cm* ⋅ *HPa*^-1^ ⋅ *d*^-1^.
  ∘ *k*_*r*_ and *k*_*x*_ of each segment calculated using the regression functions.
- *sg-sub* : an *in silico* counter-witness with no secondary growth and no suberin.
  ∘ Hydrophobic barriers fixed to *En*.*CS-Ex*.*LC* for all root segments.
  ∘ Root radius fixed to 0.198 *mm* for all root segments.
  ∘ *kAQP* fixed at 5 × 10^-4^ *cm* ⋅ *HPa*^-1^ ⋅ *d*^-1^.
  ∘ *k*_*r*_ and *k*_*x*_ of each segment calculated using the regression functions.

As radius and *k*_*x*_ regression functions are exponential, we set maximum radius at 2 mm and maximum *k*_*r*_ at 2E-10 *m*^3^*MPa*^-1^*s*^-1^.

We implemented a framework to loop the computation of root system hydraulic architecture (RSHA) for each desired timestep of each desired RSA. For each iteration, MARSHAL (Meunier et al., 2020) was run under homogeneous soil water potential, with boundary collar water potential at -15kPa. Root system hydraulic conductance *K*_*rs*_ and standart uptake fractions *SUF* were evaluated every 10 days from 10 to 60 days old RSA.

Statistics on *K*_*rs*_ and *SUF* were made using linear regressions and estimated marginal means. Normality of the data was evaluated by looking at the residuals of the linear regressions.

### Data analysis and packages

All statistical analysis were performed using R software (R Core Team, 2022).

The function ‘*lm’* from the ‘*stat’* package was used to build linear regressions. The function *‘cor’* from the ‘*stat’* package was used to compute Pearson correlation coefficients. The function ‘*shapiro*.*test’* from the ‘*stat’* package was used to perform a Sahpiro-Wilk normality test. The function ‘*boxcox’* from the package ‘*MASS’* was used to estimate *λ* parameters of Box-Cox transformations. The function ‘*emmeans*’ from ‘*emmeans*’ package was used to compute estimated marginal means. Graphs were produced using ‘*ggplot2’, ‘ggthemes’* and *‘vidiris’* packages.

### Use of artificial intelligence

Some portions of text were reformulated with AI tools, namely LeChat (https://mistral.ai/) and ChatGPT (https://openai.com/), to enhance clarity and coherence while preserving scientific consistency. Here is an example of prompt used to format results from bullet points to text : “Act as a scientist specialized in plant science. Write or rephrase what I give you in a scientific way. Be precise and use a scientific language. Avoid complex and pompous formulations. Generate only the text, nothing more.” All generated content was critically edited and reviewed.

## Abbreviations

*k*_*r*_: Radial Hydraulic Conductivity [*LT*^-1^*P*^-1^]
*K*_*r*_: Radial Hydraulic Conductance (expressed per unit of root length) [*L*^2^ *T*^-1^ *P*^-1^]
*k*_*x*_: Specific Axial Hydraulic Conductance [*L*^4^ *T*^-1^ *P*^-1^]
*kAQP*: Contribution of aquaporins to membrane hydraulic conductivity [*LT*^-1^*P*^-1^]
*SUF*: Standard Uptake Fractions [−]
*K*_*rs*_: Root System Hydraulic Conductance [*L*^3^ *T*^-1^ *P*^-1^]

## Data and Coding Availability

All codes are available here : https://doi.org/10.5281/zenodo.18324981, under Creative Common Attribution license.

## Acknowledgments

Computational resources have been provided by the supercomputing facilities of the Université catholique de Louvain (CISM/UCL) and the Consortium des Équipements de Calcul Intensif en Fédération Wallonie Bruxelles (CÉCI) funded by the Fonds de la Recherche Scientifique de Belgique (F.R.S.-FNRS) under convention 2.5020.11 and by the Walloon Region.

We are thankful to Thomas Dagbert for technical assistance in aeroponic design, and to all PEPA lab members (https://www.pepa.science/) for feedback and suggestions.

## Author contributions

**Marco D’Agostino**: Writing – Original draft, Software (GRANAR), Validation, Formal Analysis, Investigation, Data Curation, Visualization. **Rémy Schoppach**: Software (GRANAR), Formal Analysis, Validation. **Adrien Heymans**: Methodology, Software (GRANAR), Writing – Review & Editing. **Valentin Couvreur**: Methodology, Software (MECHA), Investigation, Supervision, Funding acquisition, Writing – Review & Editing. **Guillaume Lobet** : Conceptualization, Supervision, Project administration, Funding acquisition, Investigation, Writing – Review & Editing.

## Financial Support

This work has been partially supported by the Fonds National pour la Recherche Scientifique (FNRS) (M.D., grant number MIS/VA-F.4524.20), (A.H, grant number FC-31701); by the Deutsche Forschungsgemeinschaft (DFG) (M.D. and G.L., grant number EXC-2070-390732324 - PhenoRob); by the Kempe Foundation (A.H, grant number JCK-2130); and by the European Union (M.D and V.C, grant number 101043083 - ThePlantWaterPump), (G.L., grant number 101125638 – DROOGHT). Valentin Couvreur is a Research Associate of the Fonds de la Recherche Scientifique (FNRS).

## Conflicts of Interest

Conflicts of Interest: None

## Supplementals

- Supplemental 1. Data (black) and regressed (red) values of anatomical parameters with age of the root. Orange is residual deviation.
- Supplemental 2. Description of the GRANAR model.
- Supplemental 3. Parameters of CRootBox.
- Supplemental 4. Parameters of MECHA.

## Notes

### Competing Interest Statement

The authors have declared no competing interest.

