## Supplementary figures and images for "Secondary Growth and Exodermal Barriers Shape Local Root Hydraulics: Modeling Insights in Tomato"

### Supplemental 1

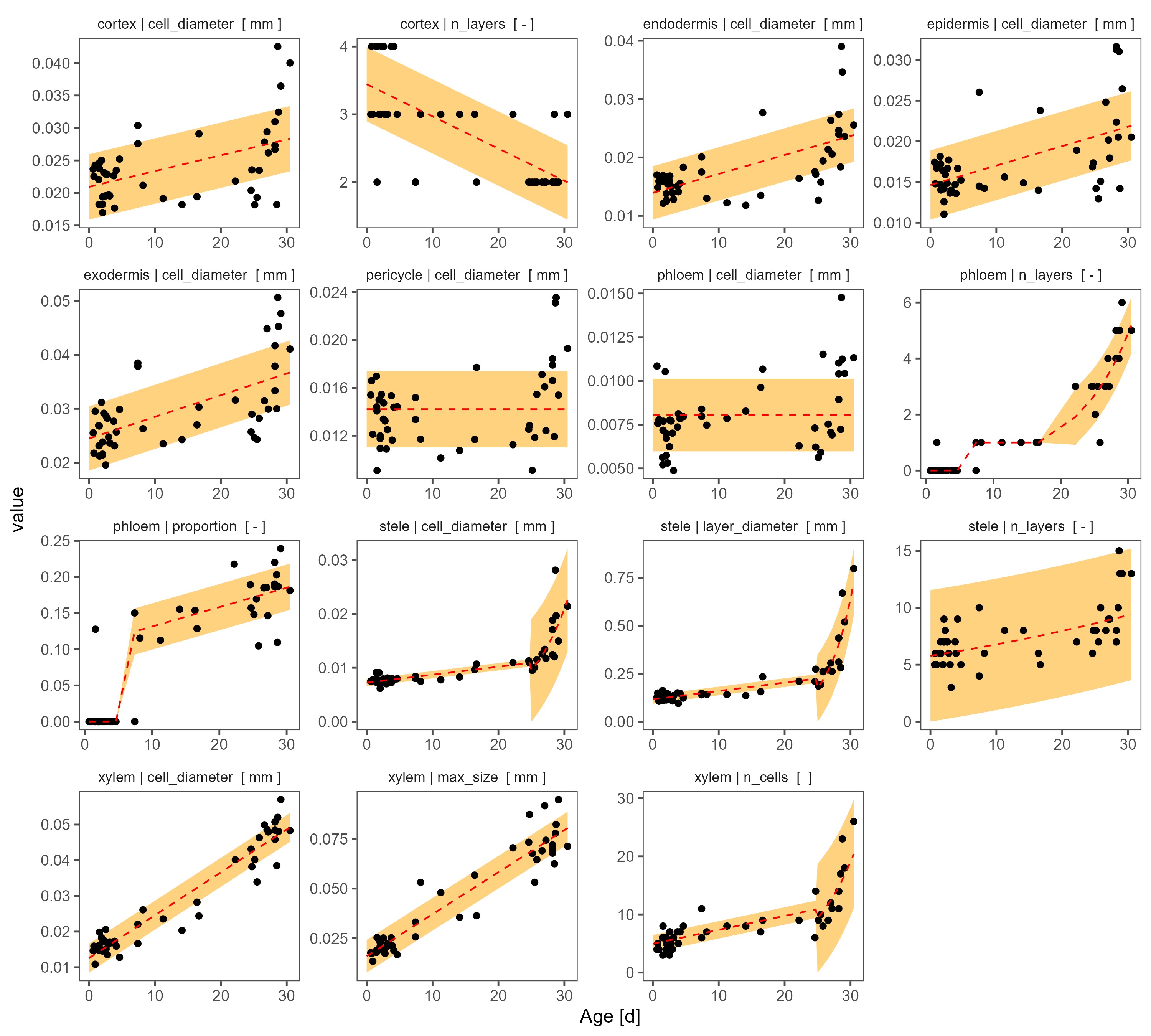
