## Supplemental 2 for "Secondary Growth and Exodermal Barriers Shape Local Root Hydraulics: Modeling Insights in Tomato"

### GRANAR : Secondary Growth Update

#### Introduction

The Generator of Root ANAtomy in R (GRANAR) is a model in R (R Core Team, 2022) to generate virtual root anatomies based on simple measurements of cross-section pictures (Heymans et al., 2020). Its purpose is to be coupled with the root hydraulic solver MECHA (Couvreur et al., 2018) to compute root radial and axial hydraulic conductivities.

This document present an update of GRANAR to generate secondary growth virtual root anatomies. All parameters are listed in Table 1.

#### Description

The code begins by creating a map of the centre of each cell, using the average size of the cell tissue and the number of layers. The process starts with the generation of the inner stele, where the total number of stele layers (which correspond to parenchyma) is determined by calculating the ratio of the total diameter of the stele to the mean diameter of the parenchyma cells. Following this, each tissue type is defined and positioned around the stele. The resulting structure consists of successive layers of tissues, beginning from the inner stele and including phloem/cambium, pericycle, endodermis, cortex, exodermis, and epidermis. For each layer of tissue, individual cells are created with a specific angle around the circular perimeter of the layer, and their corresponding coordinates relative to the centre are recorded.

Next, the inner stele is changed to add xylem, parenchyma and phloem, depending on the value of the $planttype$ parameter (which can be 1 for monocotyledons or 2 for dicotyledons) and the $secondarygrowth$ parameter (which can be 0 or 1). If $planttype$ is 1 or 2 with $secondarygrowth$ set to 0, the function *vascular()* is executed, similarly to the initial version of Heymans et al. (2020). Else, if $planttype$ is 2 and $secondarygrowth$ is 1, the new function *pack_xylem()* is executed. The workflow is the following :

1. The **number of xylem vessels** ($nC_{xylem}$) and the **mean xylem vessel diameter** ($\overline{CD_{xylem}}$) are set from data and are used to set the **xylem area** ($A_x$) :

$$A_{xylem} = nC_{xylem} \cdot \pi \left( \frac{\overline{CD_{xylem}}}{2} \right)^2 \; [mm^2]$$

2. The **stele layer diameter** ($LD_{stele}$) and the **mean stele cell diameter** ($\overline{CD_{stele}}$) are set from data, and the **mean stele cell area** ($CA_{stele}$) is calculated under the hypothesis of perfectly round cells :

$$\overline{CA_{stele}} = \pi \left( \frac{1}{2} \overline{CD_{stele}} \right)^2 \; [mm^2]$$

3. The **maximum xylem vessel area** ($maxCA_{xylem}$) is computed using the **maximum xylem vessel diameter** ($maxCD_{xylem}$).

$$maxCD_{xylem} = \pi \left( \frac{1}{2} maxCA_{xylem} \right)^2 \; [mm^2]$$

4. The **number of stele parenchyma cells** ($nC_{stele}$) is set as the difference between stele and xylem area, divided by **mean stele parenchyma cell area** ($CA_{stele}$):

$$nC_{stele} = \frac{\left((\pi \left(\frac{1}{2} LD_{stele}\right)^2 - A_{xylem}\right)}{\overline{CA_{stele}}} [-]$$

If this number is too much, it main raise errors, so we correct it a little.

5. Each stele parenchyma cell is associated with an identifier, and with an area picked in a normal distribution with $\overline{S_{cellArea}}$ as mean and a standard deviation of 10%.
6. Similarly, xylem vessels are associated with an identifier, and with an area picked in a beta distribution of parameter $\alpha = 2$ and $\beta = 4$.
7. Stele parenchyma and xylem vessel are stored in a way that xylem vessels are in the 10% most central cells.
8. The *circleProgressiveLayout* function from the *PackCircle*[1] package is applied to the stele parenchyma and xylem areas. That arranges the cells tangentially with no overlapping.
9. Xylem vessels (currently circles) are reshaped to be more realistic, by adding angles.

Eventually, the whole anatomy is generated similarly to the initial version of GRANAR. We improved and optimized other functions of GRANAR, notably the smoothing of cells and the polygon management with the *sf* Package (Pebesma, 2018). The new version of GRANAR generates cross-sections in about 1.5 seconds instead of 30 sec, no matter what type of anatomy.

[1] https://rdocumentation.org/packages/packcircles/versions/0.3.5

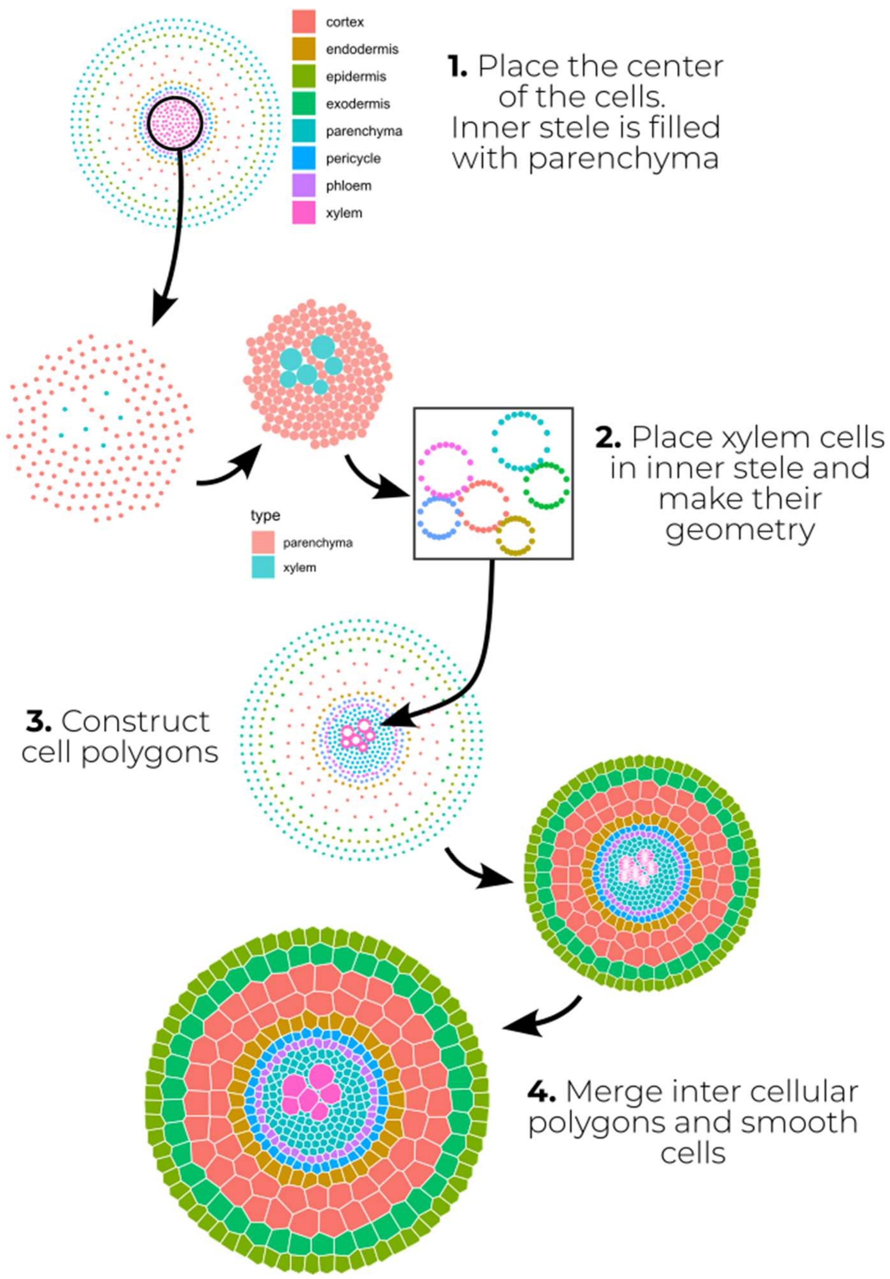


*Figure 1. Visualisation of the generation of a GRANAR anatomy.*

*Table 1. List of GRANAR parameters.*

| Name/Tissue | Type | Description | Range/units |
|---|---|---|---|
| Secondary growth | Parameter | secondary growth presence | 0 or 1 |
| Randomness | Parameter | level of randomness in the generation | 0 to 5 |
| Plant type | Parameter | plant type (monocotyledon or dicotyledon) | 1 (mono.), 2 (dico.) |
| Aerenchyma | Proportion | proportion of aerenchyma | - |
| Aerenchyma | N_files | number of aerenchyma files | - |
| Aerenchyma* | Type | type of aerenchyma : 1 for strip, 2 for patches | 1 or 2 |
| Intercellular Space* | Size | size of the intercellular space | mm |
| Intercellular Space* | Ratio | proportion of intecellular space at triple cell junctions in the cortex | - |
| Pith* | Layer_diameter | diameter of the pith layer | mm |
| Pith* | Cell_diameter | cell diameter of pith cells | mm |
| Hair* | Lengths | length of root hair | mm |
| Hair* | N_files | number of root hairs | - |
| Stele | Cell_diameter | mean diameter of inner stele (parenchyma) cells | mm |
| Stele | SD | [optional] standard deviation of inner stele (parenchyma) cell size | mm |
| Stele | Layer_diameter | diameter of the stele | mm |
| Stele | Order | ranking of the parenchyma cell layers position (usually 1) | - |
| Xylem | Max_size | maximum diameter of xylem vessels | mm |
| Xylem | Cell_diameter | mean size of xylem vessels | mm |
| Xylem | N_files | number of files (blades) of xylem (only for non-secondary growth dicot) | - |
| Xylem | N_cells | number of cells of xylem | - |
| Xylem | Order | position ranking of xylem (usually 1.5) | - |
| Phloem | N_layers | number of phloem layers (only for secondary growth dicots) | - |
| Phloem | Cell_diameter | diameter of phloem cells | mm |
| Phloem | Proportion | proportion of phloem in the stele | - |
| Phloem | Order | position ranking of phloem (usually 1.5) | - |
| Pericycle | Cell_diameter | mean pericycle cell diameter | mm |
| Pericycle | N_layers | number of pericycle layers (usually 1) | - |
| Pericycle | Order | position ranking of pericycle (usually 2) | - |
| Endodermis | Cell_diameter | mean endodermis cell diameter | mm |
| Endodermis | N_layers | number of endodermis layers (usually 1) | - |
| Endodermis | Order | position ranking of pericycle (usually 3) | - |
| Cortex | Cell_diameter | mean cortex cell diameter | mm |
| Cortex | N_layers | number of cortex layers | - |
| Cortex | Order | position ranking of the cortex layer | - |
| Inner Cortex* | Cell_diameter | mean inner cortex cell diameter | mm |
| Inner Cortex* | N_layers | number of cortex layers | - |
| Inner Cortex* | Order | position ranking of the inner cortex layer | - |
| Outer Cortex* | Cell_diameter | mean outer cell diameter | mm |
| Outer Cortex* | N_layers | number of outer cortex layers | - |
| Outer Cortex* | Order | position ranking of the outer cortex layer | - |
| Exodermis | Cell_diameter | mean exodermis cell diameter | mm |
| Exodermis | N_layers | number of exodermis layers | - |
| Exodermis | Order | position ranking of the exodermis layer | - |
| Epidermis | Cell_diameter | mean epidermis cell diameter | mm |
| Epidermis | N_layers | number of epidermis layers | - |
| Epidermis | Order | position ranking of the epidermis layer | - |
